# Auxin production in the green alga *Chlamydomonas* involves an extracellular L-amino acid oxidase and supports algal-bacterial mutualism with methylobacteria

**DOI:** 10.1101/2022.10.02.510520

**Authors:** Victoria Calatrava, Erik F. Y. Hom, Angel Llamas, Emilio Fernández, Aurora Galván

**Affiliations:** Departamento de Bioquímica y Biología Molecular. Campus de Rabanales y Campus Internacional de Excelencia Agroalimentario (CeiA3). Edificio Severo Ochoa, Universidad de Córdoba, 14071, Spain; Department of Plant Biology, Carnegie Institution for Science, Stanford, CA 94305, USA; Department of Biology and Center for Biodiversity and Conservation Research, University of Mississippi, University, MS 38677-1848, USA

**Keywords:** auxin, microalgae, mutualism, *Methylobacterium*, interkingdom signaling

## Abstract

Interactions between algae and bacteria are widespread in aquatic and terrestrial ecosystems and play fundamental roles in nutrient cycling and biomass production. However, the chemical basis for many of these interactions is poorly characterized and understood. Recent studies have shown that the plant auxin indole acetic acid (IAA) can mediate chemical crosstalk between algae and bacteria, resembling its role in plant-bacterial associations. While algae have been shown to produce IAA, molecular pathways for IAA synthesis in algae have remained elusive. Here, we report a mechanism for IAA production from L-tryptophan mediated by the extracellular enzyme L-amino acid oxidase (LAO1) in the model alga *Chlamydomonas reinhardtii*. Under inorganic nitrogen limitation but in the presence of L-tryptophan and other amino acids, high levels of IAA are generated in an LAO1-dependent manner. Thus, LAO1 plays a dual role in scavenging nitrogen from L-amino acids and in producing the phytohormone IAA, which subsequently inhibits algal cell multiplication and chlorophyll degradation. We show that these inhibitory effects can be relieved in the presence of *Methylobacterium* spp., well-known plant growth-promoting bacteria (PGPB), whose growth is mutualistically enhanced by the presence of the alga. These findings reveal a complex interplay of microbial auxin production and degradation by algal-bacterial consortia under nitrogen limitation and draws attention to potential ecophysiological roles of terrestrial microalgae and PGPB in association with land plants.

## INTRODUCTION

Photoautotrophic microalgae and cyanobacteria produce approximately half of the oxygen in the atmosphere (Field et al., 1998) and nearly half of the carbon dioxide fixed by these microbes is assimilated by heterotrophic bacteria (Fuhrman and Azam, 1982). Numerically, phytoplankton and heterotrophic bacteria dominate marine and freshwater environments (Sarmento and Gasol, 2012) and together control the nutrient cycling and biomass production at the base of the food web (Cirri and Pohnert, 2019; Rooney-Varga et al., 2005; Seymour et al., 2017). Some of these beneficial interactions have been exploited for biotechnological applications (Fakhimi et al., 2020; Wang et al., 2017) and to improve sustainable agriculture (Gonçalves, 2021; Kang et al., 2021). Nonetheless, we still understand surprisingly little about the ecology and the molecular mechanisms underlying most algal-bacterial interactions in nature (Johansson et al., 2019; Ramanan et al., 2016; Seymour et al., 2017).

There is mounting evidence highlighting conspicuous similarities between the bacterial associations formed with algae and those that form with land plants (reviewed by (Seymour et al., 2017). For instance, bacteria that associate with algae, such as *Rhizobium* and *Sphingomonas*, are often phylogenetically similar to those that form symbioses with land plants, suggesting broad eco-evolutionary affinities between bacteria and the plant kingdom (Durán et al., 2022; Krohn-Molt et al., 2017; Ramanan et al., 2015). The ‘phycosphere’, the algal analog of the plant rhizosphere, is enriched in metabolites exuded by algae that can be identical to those released by the roots of plants (Amin et al., 2015; Bell and Mitchell, 1972; Shibl et al., 2020). These compounds include nutrients such as sugars and amino acids that are exchanged with bacteria as well as signaling molecules that are essential for crosstalk between the two taxa (Segev et al., 2016).

The auxin indole-3-acetic acid (IAA) is the most extensively studied signaling molecule controlling both algal- and plant-bacterial interactions (Amin et al., 2015; Keswani et al., 2020; Lee et al., 2019; Segev et al., 2016; Tzipilevich, 2020). Since the discovery of IAA in the 1930s (Went, 1935), efforts to understand the molecular mechanisms for its synthesis and perception by plants have grown exponentially (Enders and Strader, 2015; Leyser, 2018; Weijers et al., 2018). Although initially thought to be produced exclusively by land plants, several studies have found that algae, bacteria and fungi can also produce IAA (Cox et al., 2017; Kiseleva et al., 2012; Sardar and Kempken, 2018). IAA plays a key role coordinating plant growth and development, affecting numerous processes such as tissue differentiation, cell division, lateral root formation and flowering (Cox et al., 2017; Leyser, 2018). IAA can also modulate the physiology of other organisms including algae, bacteria, fungi and animals (Amin et al., 2015; Bhoi et al., 2021; Chung et al., 2018; Fu et al., 2015; Van Puyvelde et al., 2011) making it a widespread interkingdom signaling molecule (Fu et al., 2015).

As in land plants, IAA plays a key role in controlling growth, photosynthetic activity and primary metabolism of their unicellular relatives, the microalgae (Amin et al., 2015; Chung et al., 2018; De-Bashan et al., 2008; Kozlova et al., 2017). Some bacteria can produce IAA to stimulate algal photosynthetic activity, primary metabolism and growth (Amin et al., 2015; De-Bashan et al., 2008; Meza et al., 2015; Segev et al., 2016). Auxins can also be produced by green, red and brown algae (Bogaert et al., 2019; Jacobs et al., 1985; Khasin et al., 2018; Labeeuw et al., 2016; Lau et al., 2009; Le Bail et al., 2010; Overbeek, 1940), although the molecular basis for *how* IAA is made and its potential role in mutualistic interactions remains essentially unknown (Labeeuw et al., 2016; Lin et al., 2022; Morffy and Strader, 2020).

In prior work, we found that the model alga *Chlamydomonas reinhardtii* (Salomé and Merchant, 2019), and plant growth-promoting bacteria (PGPB) in the genus *Methylobacterium* (Zhang et al., 2021) can form a mutualism based on a carbon (C) and nitrogen (N) nutrient exchange (Calatrava et al., 2018). This beneficial interaction between *Chlamydomonas* and *Methylobacterium* can be harnessed to improve wastewater bioremediation, biomass generation and hydrogen production (Torres et al., 2022). These bacteria can mineralize exogenous amino acids and peptides that are poor N sources for *Chlamydomonas* to support the growth of the alga, which in turn provisions photosynthetic C sources to promote bacterial growth (Calatrava et al., 2018, 2019). Notwithstanding, this alga can grow axenically on most free amino acids and some short peptides, relying on an extracellular L-amino acid oxidase (LAAO) named LAO1 (Calatrava et al., 2019; Muñoz-Blanco et al., 1990a; Vallon et al., 1993). LAO1 is highly expressed in *Chlamydomonas* during nitrogen starvation and oxidizes L-amino acids present in the extracellular medium to produce ammonium, hydrogen peroxide, and corresponding keto acids (Table S1). While ammonium is used to support algal growth, the resulting keto acids are paradoxically not used by the alga as a C source and remains extracellular (Chaiboonchoe et al., 2016; Muñoz-Blanco et al., 1990b; Vallon et al., 1993). We reasoned that while not metabolized by *Chlamydomonas*, these keto acids may provide some benefit and/or play some ecological role. For instance, some amino acid derived keto acids like α-ketoisocaproic (from L-leucine) or α-ketoisovaleric (from L-valine) show siderophore activity that may improve iron nutrition (Drechsel et al., 1993); others, like pyruvate (from L-alanine) and oxaloacetate (from L-aspartate), can scavenge hydrogen peroxide (Kim et al., 2015) to potentially reduce oxidative stress; still others may serve as carbon sources for nearby organisms. Some LAO1-derived keto acids like indole-pyruvic acid (IPyA, from L-Trp) and phenyl-pyruvic acid (PPA, from L-phenylalanine), are well-known precursors for auxin biosynthesis (Wang et al., 2019; Zhao, 2012). The ecological relevance of LAAO enzymes has largely been neglected, and LAAO-mediated production of auxin has only recently been identified in a single bacterial species (Rodrigues et al., 2016).

In this work, we present the first genetic evidence for algal auxin biosynthesis in the model alga *Chlamydomonas*. This occurs under N limitation in a LAO1-dependent manner, expanding the role of LAAOs in microbial auxin production. We show the impact of IAA accumulation in the extracellular medium under N limitation on *Chlamydomonas* growth and its potential role in establishing beneficial interactions with methylobacteria. Our study reveals a new mode of algal-bacterial interaction mediated by algal-produced IAA that may have fundamental implications about the ecological impact of algae and PGPB in aquatic systems and in modulating plant responses (Lee and Ryu, 2021).

## RESULTS

### *Chlamydomonas* produces IAA from tryptophan using the extracellular L-amino acid oxidase LAO1

L-tryptophan (L-Trp) is the major substrate for IAA production in bacteria, fungi, algae and land plants (reviewed by Morffy and Strader, 2020). *Chlamydomonas* can transform exogenous L-Trp and other amino acids into ammonium and a corresponding α-keto acid by means of the extracellular deaminase LAO1 (Muñoz-Blanco et al., 1990b). This enzyme is highly induced by nitrogen (N) limitation and supports growth on L-Trp as the sole N source; *lao1* knockout mutants completely lose the ability to grow under these conditions (Calatrava et al. (2019) and **Figure S1A**). To test whether *Chlamydomonas* LAO1 is involved in IAA biosynthesis, log-phase wild-type (WT) and *lao1* null mutant cells were incubated for 48 hours in growth medium supplemented with 5 mM L-Trp as a sole source of N (**Figure 1**). The concentrations of L-Trp, the corresponding keto-acid indole-3-pyruvic acid (IPyA) and IAA were quantified in cell-free supernatants (**Figure 1A, B** and **C**, respectively). A control without cells and another without L-Trp were similarly incubated (**Figure S2**). WT cultures depleted L-Trp and accumulated the products IPyA and IAA in the medium, along with other unidentified indole-containing compounds (**Figure S3**). This was not observed with the *lao1* mutant cultures or controls without cells, for which L-Trp concentration remained constant (**Figure 1A-C, Figure S3**).

**Figure 1.**
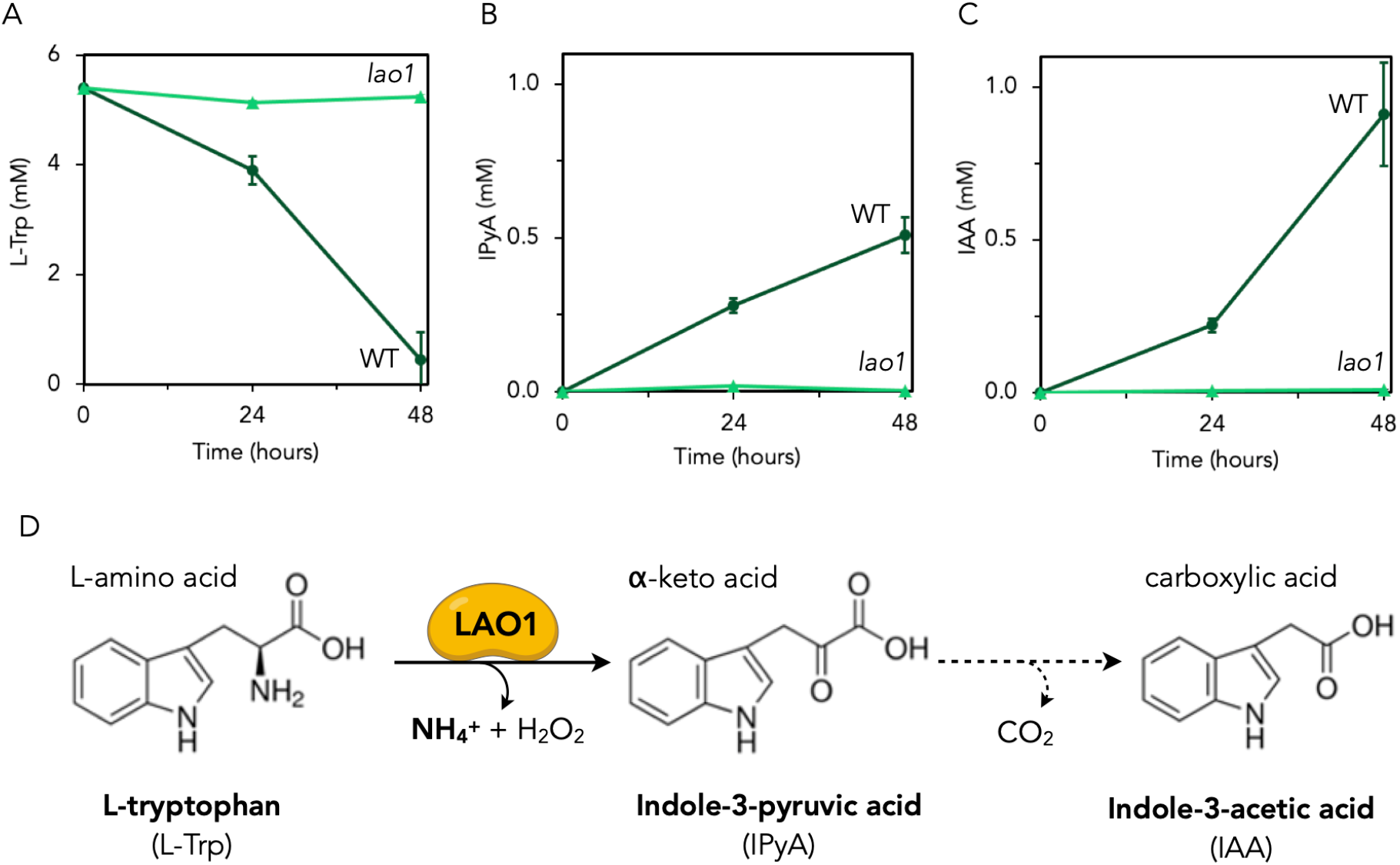
L-Amino Oxidase (LAO1) plays a critical role in indole-3-acetic acid (IAA) production from L-tryptophan (L-Trp) in *Chlamydomonas*. *Chlamydomonas* wild-type (dark green circles) and *lao1* null mutant (light green triangles) log-phase cells at 5×10^6^ cells/ml were incubated for 48 hours in nitrogen-free medium supplemented with 5 mM L-Trp as a sole source of nitrogen. (*A*) L-Trp, (*B*) indole-3-pyruvic acid (IPyA) and (*C*) IAA were quantified in the cell-free supernatants by HPLC. (*D*) LAO1 deaminates extracellular L-Trp to produce the α-keto acid, IPyA, which is decarboxylated to IAA by means of a yet unidentified mechanism (dashed arrow).

These results show that LAO1 is essential for the biosynthesis of IAA in *Chlamydomonas*. While LAO1 mediates the deamination of L-Trp into IPyA, the first step in IAA biosynthesis, the subsequent step(s) may occur spontaneously by a non-enzymatic decarboxylation of the keto acid by hydrogen peroxide (Kim et al., 2015) or may be catalyzed by an enzyme(s) (**Figure 1D)**. We were unsuccessful in generating definitive support for either possibility (**Table S2**) so the precise mechanism for IPyA conversion to IAA remains to be determined.

### IAA prevents *Chlamydomonas* from multiplying and attenuates chlorophyll degradation under N limitation

In plants, auxin biosynthesis and accumulation can be induced by nitrogen deficiency to control cell growth (Hu et al., 2020). Since we found that *Chlamydomonas* accumulates IAA under N-limiting conditions with L-Trp as the only bioavailable N source, we asked whether this auxin has a similar effect on algal growth under N limitation (i.e., in the absence of inorganic N). First, we tested the effect of L-Trp and LAO1 on *Chlamydomonas* growing on different N sources (**Figure S1**). As previously demonstrated, LAO1 was required for supporting growth on L-Ala and L-Trp as sole sources of N; growth of *lao1* mutant cells on L-Arg as a sole N source was possible because L-Arg can be transported into the cell, thus obviating the need for LAO1 activity (Calatrava et al., 2019) (**Figure S1A**). For WT cells grown on L-Arg, increasing concentrations of L-Trp had a concentration-dependent effect; at lower L-Trp concentrations (up to 2.5 mM) growth was improved, but at higher concentrations growth was inhibited (**Figure S1B**). Growth of the *lao1* mutant was not affected by L-Trp supplementation, suggesting that this concentration-dependent effect of L-Trp is due to LAO1 activity. A similar effect was observed in WT cells growing on L-Ala (**Figure S1C**), as well as in the absence of any additional N source (**Figure S1D**). No impact on growth was observed for the WT (nor for the *lao1* mutant) when grown on ammonium, which represses *LAO1* gene expression (Calatrava et al., 2019), consistent with the role of LAO1 in the concentration-dependent effect of L-Trp.

To better understand the effect of LAO1-produced IAA on *Chlamydomonas* growth, cells were supplemented with different concentrations of L-Trp, IPyA and IAA; this was conducted in the presence of L-Ala to ensure basal growth and that LAO1 is expressed (**Figure 2A-C**). A slight increase in cell yield was observed at 1 mM L-Trp. This could be explained by a slightly higher total N input and/or due to the production of IAA, which could have a stimulatory effect at low concentrations (Park et al., 2013). Growth reduction was observed at higher concentrations of L-Trp, however, which could be attributed to the accumulation of IAA and other byproducts. Indeed, IAA concentrations higher than 10 mM had a clear inhibitory effect on culture cell density, with almost 70% inhibition at 100 mM IAA (**Figure 2C**). In contrast, similar levels of the intermediate IPyA did not affect growth (**Figure 2B**), consistent with the inability we observed of producing IAA directly from IPyA under these conditions (**Table S2**).

**Figure 2.**
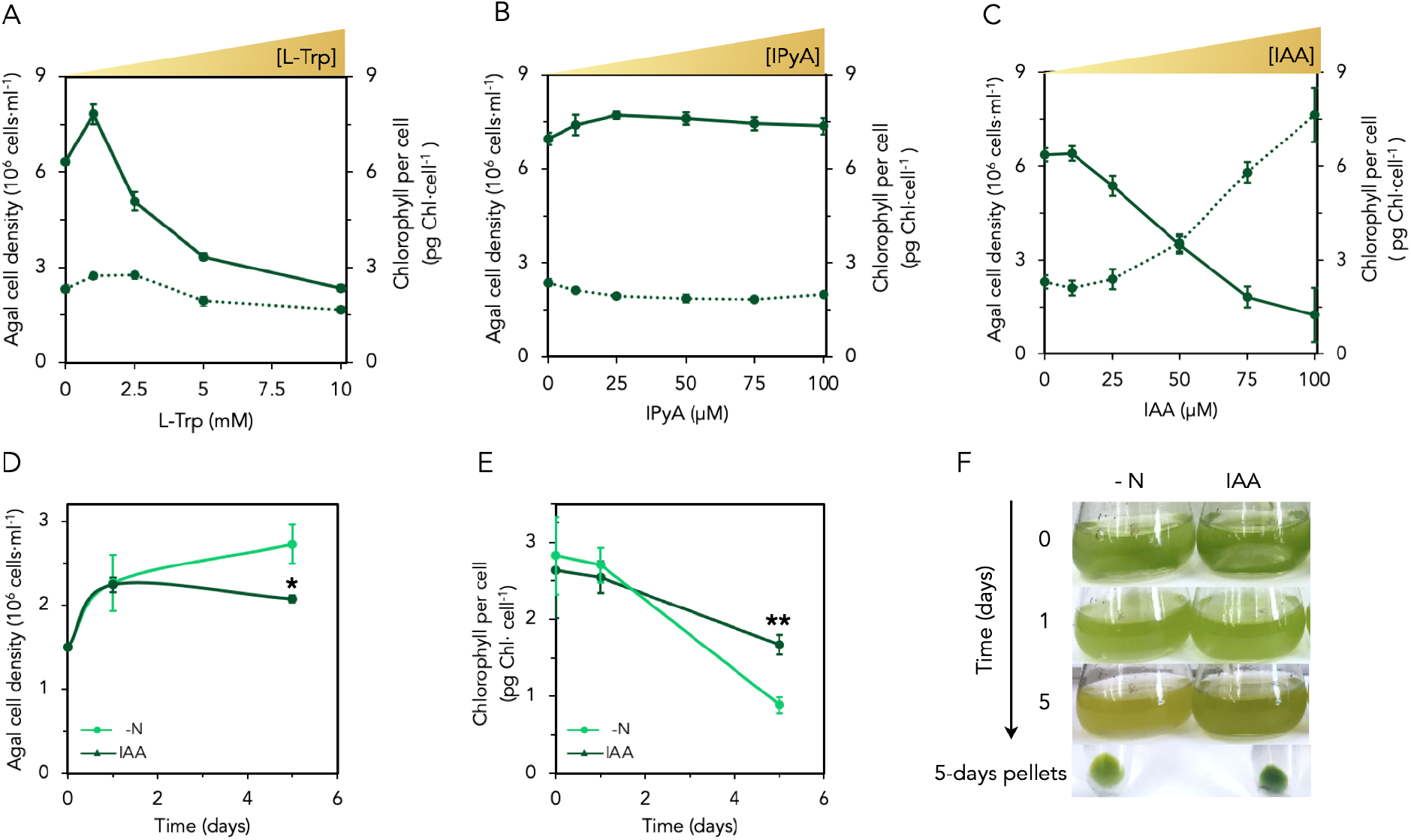
IAA arrests cell multiplication and attenuates chlorophyll degradation in nitrogen starved *Chlamydomonas*. (*A-C*) Impact of exogenously added tryptophan (L-Trp), indole-3-pyruvic acid (IPyA) and indole-3-acetic acid (IAA) on *Chlamydomonas* growth in the presence of L-alanine (L-Ala). Wild-type cells at an initial concentration of 0.2×10^6^ cells/ml were grown for three days on L-Ala (4 mM) as a N source (to enable algal growth under N-limiting conditions and ensuring LAO1 expression; see *Methods*) and supplemented with (*A*) L-Trp, (*B*) IPyA or (*C*) IAA. Cell culture densities are indicated as solid lines and chlorophyll content per cell as dotted lines. (*D-F*) Impact of IAA in *Chlamydomonas* during N deprivation. (*D*) Cell density and (*E*) chlorophyll content during N deprivation (in the absence of any assimilable N source). Wild-type cells were incubated in N-free media (–N) or supplemented with 500 *µ*M of IAA (IAA). Data are averages of 3 biological replicates with error bars depicting standard deviations. Asterisks indicate statistically significant differences compared to the control without IAA (t-test: n=3;α=0.05). (*F*) A representative culture flask of each condition in panels *D-E* was imaged at the start, after one day, and after five days; cell pellets shown were harvested by centrifugation after five days of incubation.

The reduced cell density observed with increasing concentrations of extracellular IAA was correlated with an increase (up to four times) in chlorophyll content per cell (**Figure 2C)**, which was not observed with IPyA **(Figure 2B**). This effect was also not observed with high levels of L-Trp (**Figure 2A**), possibly because additional intermediates produced from L-Trp deamination (e.g., hydrogen peroxide) may impair the accumulation of chlorophyll. N limitation generally leads to chlorophyll degradation (Schmollinger et al., 2014), so the higher chlorophyll content per cell we observe in response to exogenous IAA could be explained by a reduction in chlorophyll degradation, which must be IAA-specific and not merely LAO1-specific. To test this, we incubated concentrated, N-starved *Chlamydomonas* cultures with IAA and indeed found that the degradation of chlorophyll during N deprivation was reduced (**Figure 2D-F, S4**). In *Chlamydomonas*, chlorophyll degradation under N deprivation is linked to the mobilization of stored N to allow cells to duplicate one additional round before the complete cessation of growth (Schmollinger et al., 2014). Here, we observed that despite being N deprived, *Chlamydomonas* cultures significantly reduce cell multiplication rate in the presence of IAA, which may explain the increased chlorophyll content of these cells.

### IAA production in *Chlamydomonas* facilitates a mutualistic interaction with methylobacteria

Since IAA mediates plant/algal-bacterial interactions, we asked whether IAA and LAO1 could play a role in the establishment of interactions of *Chlamydomonas* with the bacterial genus of *Methylobacterium* given prior work demonstrating mutualistic tendencies of this taxa with *Chlamydomonas* under N-limiting conditions (Calatrava et al., 2018). Following up on this work, we quantified the growth of *Chlamydomonas* in co-culture with nine different species of methylobacteria using L-Trp as the sole nitrogen source. We observed that co-culturing with different *Methylobacterium* spp., including *M. aquaticum*, improved algal growth (**Figure S5**). Naively, this growth promotion could simply be due to methylobacteria mineralizing L-Trp to provision *Chlamydomonas* with ammonium, which is more efficiently used by this alga than L-Trp as we similarly reported for L-proline (Calatrava et al., 2018). However, this seems unlikely because we observed no algal growth promotion of the *lao1* knockout mutant by methylobacteria (**Figure S5**), suggesting that LAO1 is essential for this growth enhancement. Given the conversion of L-Trp to IAA and the inhibitory effects of IAA on *Chlamydomonas*, we hypothesized that methylobacteria may promote algal cell growth by reducing the levels of IAA accumulated in the media in co-culture relative to *Chlamydomonas* in monoculture. Indeed, in co-culturing with *M. aquaticum*, IAA-induced inhibition of cell multiplication and chlorophyll degradation was reduced to levels similar to conditions without IAA (**Figure 3B**,**C**) and measured auxin levels were highly reduced (**Figure 3D**), strongly supporting a methylobacteria-mediated degradation of IAA. Since auxin concentrations were not reduced in bacterial monocultures nor in algal monocultures, both microbes are necessary for auxin to be eliminated from the media in a cooperative fashion. This effect was observed with nine other *Methylobacterium* spp. although was most pronounced with *M. aquaticum* (**Figure 3A)**. Importantly, co-culturing with exogenously added IAA resulted in a mutualistic increase in cell density for both microbes compared to the respective monocultures (**Figure 3E-G**); *M. aquaticum* increased its cell density ~60-fold whereas *Chlamydomonas* increased by ~1.5 times in IAA-supplemented co-culture (**Figure 3E**). Since no N source other than IAA was added to the media, this observation strongly suggests that the bacterium, prompted by the presence of the alga, is feeding on IAA as a N (and likely carbon) source. On the other hand, since no detectable ammonium was produced by the bacterial monocultures on IAA (**Table S3**), *Chlamydomonas* most likely benefits from relief of IAA-induced inhibition through the bacterial degradation of this auxin.

**Figure 3.**
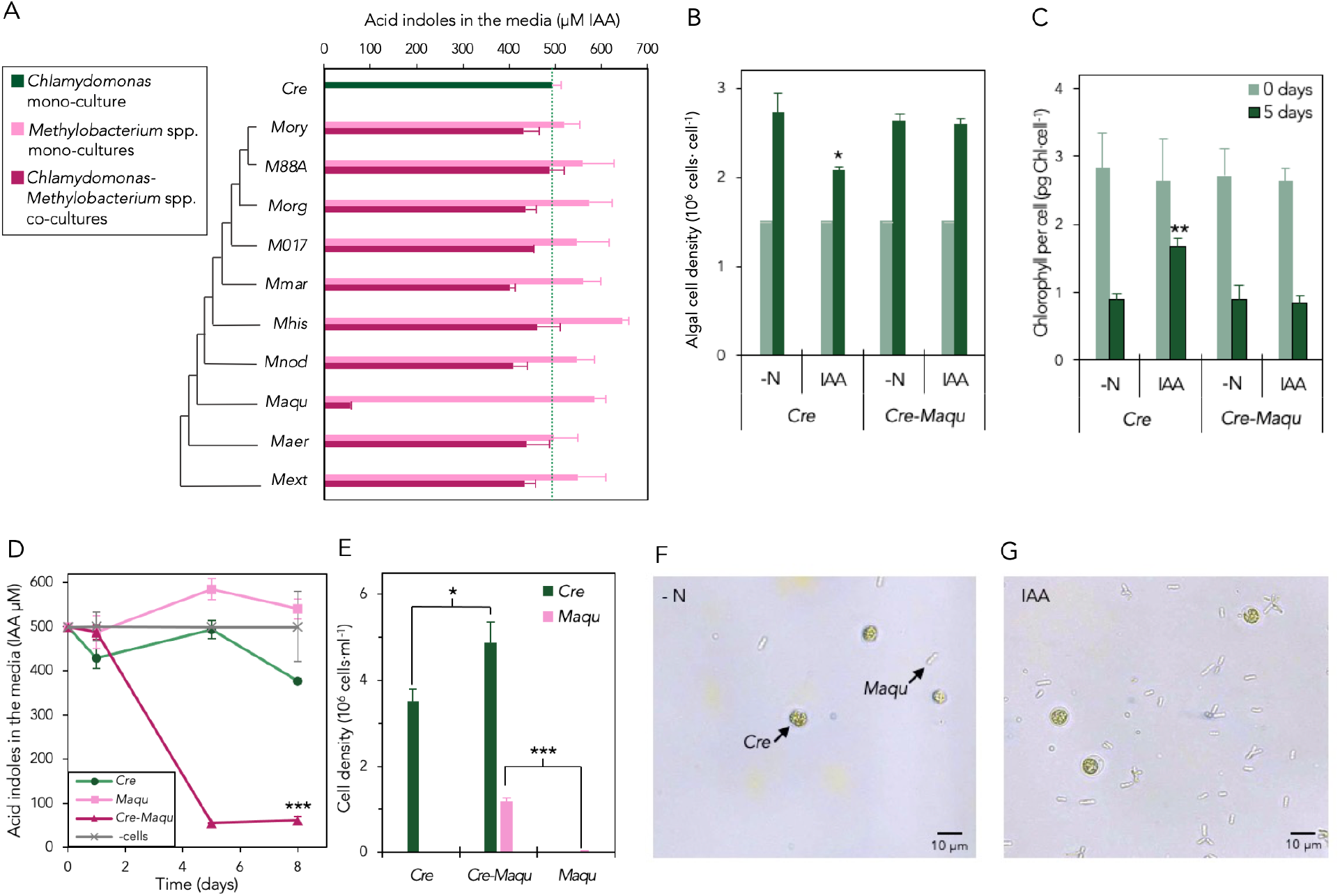
*Methylobacterium* spp. reduces IAA levels in *Chlamydomonas* cultures to relieve algal inhibition of cell multiplication and chlorophyll degradation. (*A*) Algal and bacterial mono- and co-cultures were incubated on N-free media supplemented with 500 *µ*M of IAA for five days. Indole concentration in the cell-free media was determined using the Salkowski reagent (see *Materials and Methods*). *Methylobacterium* spp. examined: *M. oryzae (Mory); M*. sp. 88A *(M88A); M. organophilum (Morg); M*. sp. M017 *(M017)*; *M. marchantiae (Mmar); M. hispanicum (Mhis); M. nodulans (Mnod); M. aquaticum (Maqu); M. aerolatum (Maer); Methylovorus extorquens* (previously known as *Methylobacterium extorquens; Mext*). Initial cell concentrations were 10^6^ cells/ml for *Chlamydomonas* and A_600_ of 0.01 for *M. aquaticum* (approximately 10_6_ cells/ml). (*B*) Algal cell density and (*C*) chlorophyll content per cell were determined initially (0 days) and after five days of incubation for *Chlamydomonas* monocultures (Cre) and *Chlamydomonas-M. aquaticum* co-cultures (Cre-Maqu) in N-free medium (–N) supplemented with 500 *µ*M IAA (IAA). (*D*) *Chlamydomonas* (Cre) monocultures, *M. aquaticum* (Maqu) monocultures and co-cultures (Cre-Maqu) were processed as in (*A*) in media supplemented with 500 *µ*M of IAA. (*E*) Algal and bacterial cell densities after eight days of growth on 500 *µ*M of IAA in co-culture or monoculture were quantified using qPCR of single-copy genes specific for *Chlamydomonas* (centrin) or *M. aquaticum* (*rpoB*) (see *Materials and Methods*). (*F*) *Chlamydomonas-M. aquaticum* co-cultures were imaged using a light microscope under conditions without a nitrogen source and (*G*) supplemented with 500 *µ*M IAA after five days. Data shown are averages of 3 biological replicates with error bars depicting standard deviations. Asterisks indicate statistically significant differences compared to the control comparison (t-test: n=3;α=0.05).

## DISCUSSION

The interactions between algae and heterotrophic bacteria play a fundamental ecological role controlling nutrient cycling and biomass production in their habitats (Cirri and Pohnert, 2019). Recent studies have shown that bacteria can produce the phytohormone IAA to promote algal growth (Amin et al., 2015), resembling the role of this molecule in plant-bacterial associations. However, auxin production by the algal partner has been largely overlooked or neglected (Amin et al., 2015; Segev et al., 2016) despite the important ecological and economic impact this may cause in both aquatic and terrestrial ecosystems (Keswani et al., 2020; Yang et al., 2017). Moreover, there has thus far been no known genetic pathway by which to produce IAA in algae. Here, we show a novel and unexpected mechanism of IAA production in the model alga *Chlamydomonas* under N-limiting conditions.

*Chlamydomonas* can deaminate L-Trp extracellularly via the LAAO enzyme LAO1 to yield the keto acid IPyA, the major precursor for auxin biosynthesis in plants, bacteria and fungi (Morffy and Strader, 2020). A pathway for tryptophan-dependent IAA production was recently discovered in the plant-benefiting bacterium *Gluconacetobacter diazotrophicus* encoded by a gene cluster containing L-amino acid oxidase, cytochrome C and ridA genes (Rodrigues et al., 2016). *Chlamydomonas* harbors a similar gene cluster that includes L-amino acid oxidase (*LAO1*) and *RidA* genes; LAO1 homologs are also found in other algae (Calatrava et al., 2019). We recently characterized a *Chlamydomonas lao1* knockout mutant that lacks extracellular LAAO activity (Calatrava et al., 2019) that we used here to evaluate the relevance of this enzyme for IAA production. We have shown that LAO1 is essential for the production of IAA from L-Trp via an IPyA intermediate. Whereas this enzyme is most likely involved in the initial step of L-Trp deamination to yield IPyA, the conversion of IPyA into IAA may be achieved by a two-step pathway mediated by ipdC decarboxylase (via indole-3-acetaldehyde intermediate) in some bacteria or by the single-step YUCCA pathway (flavin monooxygenase-like enzyme) in plants (Morffy and Strader, 2020). As in *G. diazotrophicus*, no homologs for any of these enzymes have been identified yet in *Chlamydomonas* and thus, the nature of the conversion of IPyA into IAA awaits experimental confirmation.

Previously limited to a single bacterial species, the role of LAAO enzymes in IAA production has been broadened here to include the microalga *Chlamydomonas* and may extend to other algae that harbor LAO1 homologs including members of Rhodophyta, Alveolata, Haptophyta, Heterokonta and Dinophyta (Calatrava et al., 2019). In support of this idea, the haptophyte alga *Emiliania huxleyi* harbors two putative *LAO1* gene homologs (Calatrava et al., 2019) and shows a ratio of L-Trp conversion to IAA similar to what we found here in *Chlamydomonas* (16-20% of L-Trp conversion) (Labeeuw et al., 2016). However, based on existing genomic data, the *LAO1* gene seems to be absent in most members of the green lineage (i.e., green algae and land plants) other than *Chlamydomonas* spp. (Calatrava et al., 2019), so they must rely on alternative pathways for IAA biosynthesis. Higher levels for IAA production in *Chlamydomonas* have been reported relative to 23 other green algal species (Stirk et al., 2013), consistent with *Chlamydomonas* producing IAA using a different strategy compared to sister lineages. To date, transcriptomic and/or bioinformatic analyses appear to indicate that the most common putative pathway for IAA biosynthesis in microalgae, including *Chlamydomonas*, occurs via an indole-3-acetamide intermediate (reviewed by Zhang et al., 2022). While we have shown that LAO1 is crucial for IAA production under N limitation, alternative pathways that remain to be elucidated may be more relevant under different conditions.

The presence of LAO1 in *Chlamydomonas* was likely acquired through horizontal gene transfer, suggesting an adaptive advantage of having LAO1 in *Chlamydomonas’* native environment (Calatrava et al., 2019). Originally isolated from soil, this alga and has been readily found in the rhizosphere (Elizabeth H. Harris, 2009) where inorganic N may be limiting but free amino acids may be present from plant exudates and microbial decay (Jaeger et al., 1999; Kravchenko et al., 2004; Rybicka, 1980). In this context, LAO1 could play a prominent role in scavenging N from a broad range of amino acids (Calatrava et al., 2019). As a periplasmically targeted enzyme in *Chlamydomonas* (Muñoz-Blanco et al., 1990a; Vallon et al., 1993), LAO1-driven deamination of amino acids may release ammonium not only for its own use but potentially as a chemoattractant (Chet and Mitchell, 1976) and/or a diffusible ‘public good’ for closely associated or nearby partners (cf. Bachmann et al., 2013; Lindsay et al., 2019; Nadell et al., 2016). The accumulation of LAO1 keto acid byproducts as public goods in the phycosphere could have additional roles that have been thus far neglected. Here, we have shown that LAO1 leads to IAA accumulation in the extracellular medium. While relatively low concentrations of IAA can improve algal growth (Park et al., 2013), we observed that the accumulation of high levels of IAA in the media arrests algal cell proliferation and chlorophyll degradation. This is consistent with previous observations of higher chlorophyll accumulation in response to IAA in the microalgae *Chlorella vulgaris, Nannochloropsis oculata* and *Acutodesmus obliquus* (Piotrowska-Niczyporuk et al., 2018; Piotrowska-Niczyporuk and Bajguz, 2014; Trinh et al., 2017). An increase in photosynthetic performance by IAA has also been observed in the microalgae *Emiliania huxleyi* and diatoms that harbor IAA-producing symbiotic bacteria (Amin et al., 2015; Labeeuw et al., 2016). Similar effects have been observed in the chloroplasts of plants (Misra and Biswal, 1980). This effect of IAA on the chlorophyll content is stronger in aged algal cultures and plant samples (Evans, 1985; Labeeuw et al., 2016; Misra and Biswal, 1980) and might be linked to a condition of stress (Gururani et al., 2015). We considered the effect of IAA on chlorophyll levels in the context of N limitation as a stress condition that triggers LAO1 accumulation. Under mixotrophic conditions (i.e., in the presence of acetate), this alga prioritizes cellular respiration over photosynthesis (Schmollinger et al., 2014). As a result, when N is limiting, pigments and other N-containing macromolecules involved in photosynthesis and chloroplast function are degraded first, presumably to enable *Chlamydomonas* cells to undergo an additional round of duplication (Schmollinger et al., 2014). The accumulation of exogenous IAA by *Chlamydomonas* could function as a quorum sensing-like signal molecule (Khasin et al., 2018; Segev et al., 2016) to decrease growth when the cell population is high and nutrient resources are limited. Preventing cell multiplication while avoiding the breakdown of the photosynthetic machinery during N limitation, may be a hedge-betting strategy to capitalize on mutualistic cross-feeding interactions with other N-mineralizing microbes in the vicinity (e.g., *Methylobacterium*). These neighboring microbes may benefit from the provisioning of photosynthates like glycerol in exchange for more readily assimilable sources of N like ammonium that *Chlamydomonas* can use to resume growth (Calatrava et al., 2018; Ietswaart et al., 1994).

IAA can be metabolized by rhizospheric bacteria (Donoso et al., 2017; Ebenau-Jehle et al., 2012; Lin et al., 2012; Scott et al., 2013), some of which like *Pseudomonas putida* show chemotaxis towards the auxin (Rico-Jiménez et al., 2022; Scott et al., 2013). Thus, IAA production by *Chlamydomonas* may attract and feed bacteria that could potentially be beneficially self-serving. When amino acids are present but inorganic N is absent, *Chlamydomonas* interacts mutualistically via C and N cross-feeding with *Methylobacterium* spp. (Calatrava et al., 2018), species that have been shown to co-occur with *Chlamydomonas* and that are part of the phycosphere of other green algae (Krug et al., 2020; Macey et al., 2020; Madhaiyan et al., 2007). Here, we demonstrate another mutualistic mode of interaction between *Chlamydomonas* and *Methylobacterium* spp., mediated through IAA. *M. aquaticum*, induced or complemented by the presence of *Chlamydomonas*, can feed on IAA, which can “release the brake” on *Chlamydomonas* growth and facilitate the enhanced growth of both organisms. From the algal perspective, LAO1-mediated production of extracellular ammonium and keto acids may thus constitute a multifaceted strategy of signaling, waiting for partner feedback, followed by mutualistic stimulation that may mirror a sort of call-and-response dynamic of plant-microbe interactions (Clear and Hom, 2019).

To our knowledge, this is the first study to report an algal-dependent mechanism for IAA degradation in bacteria and IAA degradation by *Methylobacterium* spp. in general. IAA catabolism (*iac*) genes (Laird et al., 2020) appear to be absent from available *Methylobacterium* genomes but we believe that IAA degradation by *M. aquaticum* (involving a yet unidentified pathway) may cause a carbon/nitrogen metabolic imbalance (Bren et al., 2016) that hampers its growth in the absence of the alga. During co-culture, algal-derived photosynthates may restore carbon/nitrogen balance and enhance bacterial growth (Calatrava et al., 2018). We cannot rule out the possibility that other putative metabolites or proteins produced by the alga may also potentially facilitate bacterial degradation of IAA. Regardless, IAA-degrading bacteria can influence not only land plants but also algae (Laird et al., 2020). A previous study found that the haptophyte microalga *Emiliania huxleyi* exhibits strain-specific differences in the production of IAA and susceptibility to infection by the pathogenic roseobacter *Ruegeria* sp. R11: while the non-IAA producer strain is resistant to roseobacter, the IAA-producing strain is susceptible to bacterial infection (Labeeuw et al., 2016). This bacterial infection is accelerated by the addition of L-Trp to the co-culture, which suggests that L-Trp-derived IAA production by the alga could enhance bacterial infection. However, since the levels of IAA in the algal cultures were shown to be reduced in co-culture with roseobacter, the role of algal-derived IAA production was not pursued (Labeeuw et al., 2016). Given the findings presented here showing that bacterial degradation of IAA can be triggered in the presence of algae, we speculate that in the presence of *E. huxleyi, Ruegeria* sp. may metabolize algal-derived IAA to achieve higher bacterial densities and potentially accelerate algal infection. Regardless, we believe that algal-dependent bacterial degradation of IAA may be relevant for many other bacterial species and their interactions with both algae and land plants.

Among the 9 *Methylobacterium* spp. tested in this work, *M. aquaticum* was significantly more efficient in degrading IAA in the presence of *Chlamydomonas*. This may underlie the basis for this particular bacterial species’ broad ability to enhance the growth of early diverging plant lineages like green algae and moss, (Calatrava et al., 2018; Tani et al., 2012). *Chlamydomonas* and *M. aquaticum* have both been found in freshwater environments and in the rhizosphere (Mendes et al., 2013; Stringlis et al., 2018; Tani et al., 2015). Given niche overlap of algae and higher plants in soil (Mendes et al., 2013; Stringlis et al., 2018) and the ability of algae to “dialogue” with other microbes using the same auxin compounds that regulate land plants growth and development, we hypothesize that algae may play an underappreciated role in the recruitment of PGPB to the plant microbiome, let alone in modulating plant fitness through algal-plant interactions (**Figure 4**). Plant-plant interactions dramatically shape plant fitness, coexistence, life histories, and community assembly (Subrahmaniam et al., 2018; Wang et al., 2021); it is plausible that interactions of plants with earlier-diverging green lineages are also important and influential (Gavini et al., 2019; Gornall et al., 2011).

**Figure 4.**
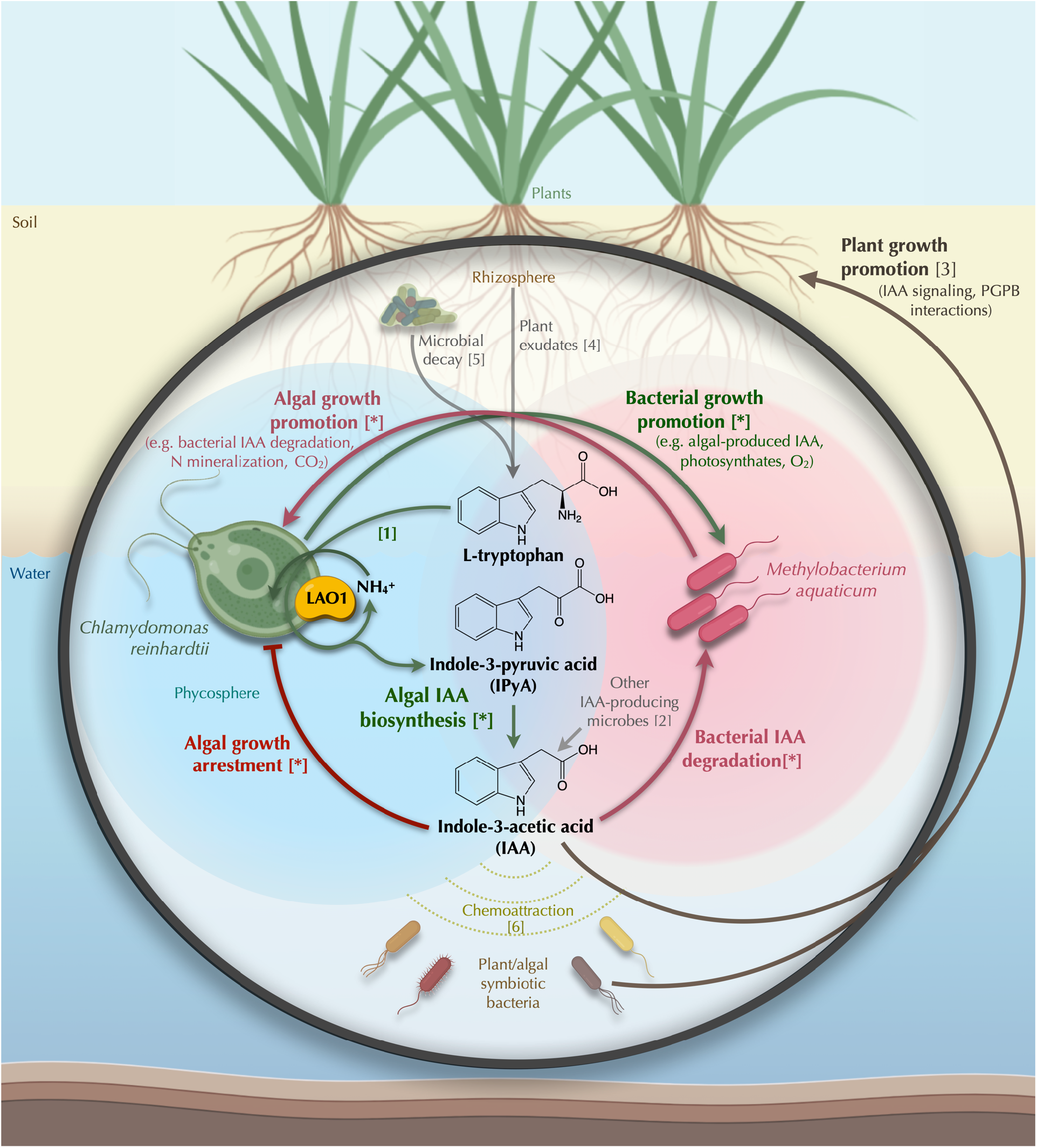
Proposed role of auxin-mediated mutualistic interactions between *Chlamydomonas, Methylobacterium* and land plants. In soil environments where tryptophan may be present due to plant exudation and microbial decay, bacterial indole-3-acetic acid (IAA) biosynthesis from this amino acid can promote plant growth. *Chlamydomonas reinhardtii* can also convert tryptophan into IAA using the extracellular enzyme LAO1. Accumulation of this auxin may result in algal growth arrest and in attracting beneficial PGPB bacteria. In the presence of the IAA-degrading bacterium *Methylobacterium aquaticum*, IAA is depleted, enhancing growth of both microorganisms. Metabolites exchanged between bacteria and algae could strengthen this mutualistic interaction (Calatrava et al., 2018, 2019). Moreover, we imagine three-way algal-plant-bacterial associations whereby algal-derived IAA not only benefits the alga but promotes plant-bacterial symbioses and modulates plant physiology directly. Figure created using BioRender and ChemDraw 20.1. [*] indicates interactions or processes shown in this work. Numbers in square brackets correspond to references: [1] (Vallon et al., 1993); [2] (Cox et al., 2017; Kiseleva et al., 2012; Sardar and Kempken, 2018); [3] (Scott et al., 2013); [4] (Kravchenko et al., 2004); [5] (Moe, 2013); [6] (Backer et al., 2018).

The role of microalgae as players in the plant microbiome has only recently started to be appreciated and their use to improve soil fertility, water preservation and plant growth is now emerging as a promising approach for sustainable agriculture (Feng et al., 2022; Gitau et al., 2022; Huo et al., 2020; Keswani et al., 2020; Lee and Ryu, 2021; Srivastava et al., 2018; Upadhyay et al., 2016); see Alvarez et al., 2021 for a recent review). For these new applications, understanding how and under what environmental conditions IAA is produced by rhizospheric and phyllospheric algae is of great interest. Bacterial degradation of IAA has been shown to be essential for rhizosphere colonization and plant growth promotion (Zúñiga et al., 2013), and similar to how LAAO-mediated auxin production is involved in plant growth promotion by *G. diazotrophicus* (Rodrigues et al., 2016), auxin production by other LAAO-containing microbes in the rhizosphere like *Chlamydomonas* could have an impact on plant growth and fitness. Our findings suggest that future studies should carefully consider the role that terrestrial algae play in plant microbiomes (Lee and Ryu, 2021) and in the historical record of land plant evolution (Puginier et al., 2022).

## CONCLUSION

We provide genetic evidence that supports IAA production by the model alga *Chlamydomonas* via an unexpected LAAO-mediated pathway, which has thus far been restricted to a single bacterial species (Rodrigues et al., 2016). This finding supports a new role for microbial LAAOs in auxin production that may be widespread in other algal lineages (Calatrava et al., 2019; Labeeuw et al., 2016). We also show that IAA may act as an extracellular self-regulatory/quorum sensing-like molecule to control cell multiplication and delay the breakdown of the photosynthetic machinery under inorganic N-limitation. This may be an ecological strategy to facilitate interactions with N-mineralizing bacteria. We demonstrate that the plant- and algal-benefiting bacteria *Methylobacterium* spp. can degrade extracellular levels of IAA generated by the alga in the presence of L-Trp and N limitation to alleviate inhibition of algal cell multiplication. Interestingly, these bacteria can feed on IAA only in the presence of the alga, revealing a new cooperative mode of auxin degradation that is induced and/or functionally complemented by the alga. This new mode suggests a role for algal-driven IAA production prompted by algal-bacterial interactions that may be relevant to plant-microbiome dynamics. Since *Chlamydomonas* and *Methylobacterium* are naturally found in the rhizosphere, their role in modulating IAA levels may impact plant fitness and could be exploited for crop improvement in sustainable agriculture. Overall, this work extends our understanding of auxin production and degradation by algae and algal-bacterial consortia under N limitation, highlighting the potential for tri-partite interactions between rhizospheric algae, PGPB, and land plants.

## MATERIAL AND METHODS

### Microbial strains and culture media

*Chlamydomonas reinhardtii* wild-type strain CC-5325 and *lao1* mutant LMJ.RY0402.044073 were obtained from the *Chlamydomonas Resource Center* (https://www.chlamycollection.org/). This *lao1* mutant contains a CIB gene cassette insertion at the *Cre12*.*g551352* locus, corresponding to the *LAO1* gene (Li et al., 2016), that prevents LAO1 function (Calatrava et al., 2019). Cells were pre-cultured for 2-3 days in Tris-Acetate-Phosphate (TAP) medium (Gorman and Levine, 1965) containing 8 mM of ammonium chloride at 23 ºC under continuous light and agitation (120 rpm).

*Methylobacterium* strains used in this work are summarized in Table 1. Cells were pre-cultured in *<u>Me</u>thylobacterium* <u>M</u>edium (MeM) as previously described (Calatrava et al., 2018) supplemented with 2.5 g·L^−1^ of tryptone (MeM^T^) for 2-3 days at 28 ºC and continuous agitation (150 rpm).

**Table 1.**
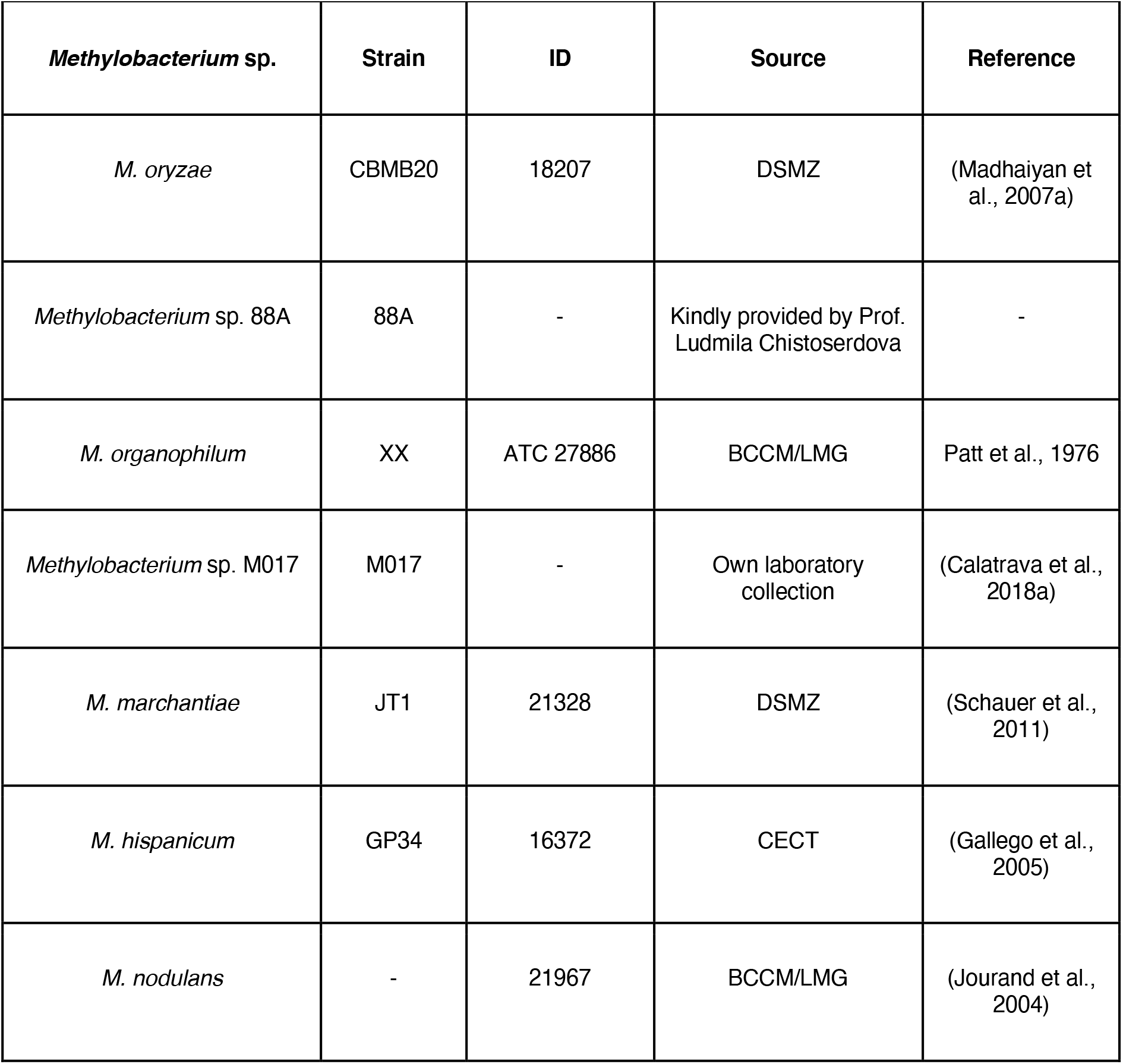
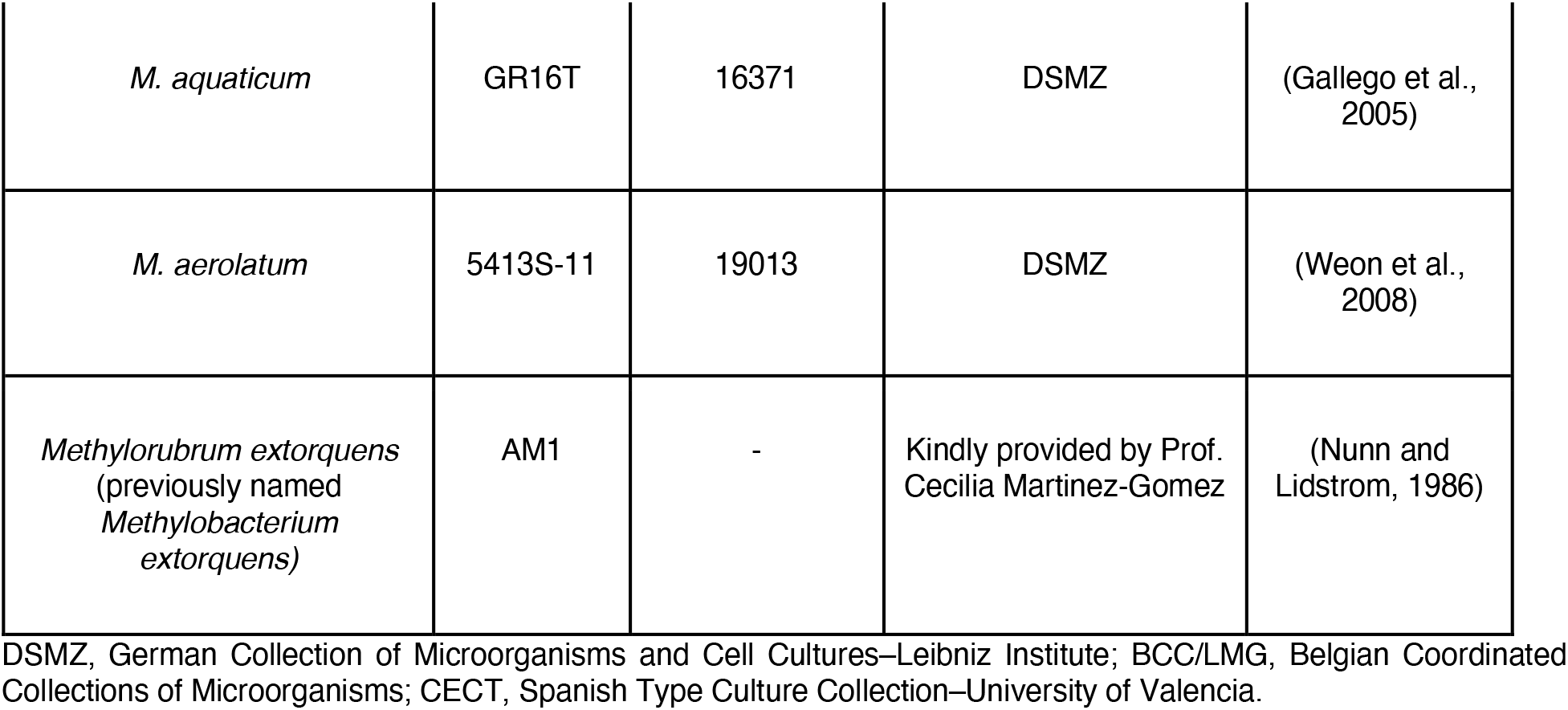
*Methylobacterium* spp. strains used in this work.

To detect potential contamination, cell inocula were routinely checked by streaking on agar plates of TAP supplemented with yeast extract (2.5 g·L^−1^), incubated for 2 weeks at 23ºC and examined under a light microscope.

### Determination of L-tryptophan metabolization, and indole-3-pyruvic acid and indole-3-acetic acid biosynthesis by *Chlamydomonas*

*Chlamydomonas* wild-type and *lao1* mutant were pre-cultured as described above to exponential phase. Cells were harvested by centrifugation for 2-3 min at 2,090 *g* and washed three times with nitrogen-free TAP medium (T-N). Cells were subsequently incubated in T-N medium supplemented with 5 mM of L-Trp under continuous light and agitation (120 rpm). After 24 and 48 hours, cell-free supernatants were stored at −20ºC until analyzed. L-Trp, indole-3-pyruvic acid and indole-3-acetic acid analysis was performed by the Chromatography Department staff at the Central Service for Research Support (SCAI) of the University of Córdoba, using High-Performance Liquid Chromatography (HPLC) and UV-Vis detection. A mixture of L-Trp (Sigma-Aldrich, Spain), indole-3-pyruvic acid (Sigma, Spain) and indole-3-acetic acid (GoldBio, US) was included as standard.

### *Chlamydomonas* cell growth tests

To test *Chlamydomonas* growth, cells were pre-cultured and washed as described above, placed in wells of a 48-well culture plate (BRANDplates, BrandTech Scientific, US) and incubated in fresh culture medium at an initial concentration of 0.2×10^6^ cells/ml. In these growth assays L-alanine was supplemented in the medium as indicated in the figure captions to allow algal growth while ensuring N-limiting conditions and LAO1 activity. At the indicated times, cell concentrations were determined using 100-200 µl in a microcell counter (Sysmex F-500, Sysmed Inc, Europe).

### Chlorophyll determination

Chlorophyll (a+b) was extracted from 1 ml of algal or algal-bacterial pellets with ethanol (1 ml) and measured spectrophotometrically (Beckman Coulter, DU 800) (Wintermans and De Mots, 1965). Chlorophyll concentration in the cultures (µg/ml) was inferred to cell concentration (cells/ml) to determine the chlorophyll content per cell.

### Indole-3-acetic acid degradation tests

To determine microbial indole-3-acetic acid depletion from the medium, *Chlamydomonas* and *Methylobacterium* spp. mono- and co-cultures were initially pre-cultured independently as described above. After 2 days (exponential growth phase), cells were washed using T-N medium and incubated in fresh T-N medium supplemented with 500 *µ*M of IAA under continuous light and agitation (120 rpm). Initial cell densities if algae and bacteria were set at 1.5×10^6^ cells/ml and A_600_=0.01 (approximately 10^6^ cells/ml), respectively. A negative control without inoculum was incubated to account for any potential abiotic degradation of IAA. After 1 week of incubation, the cell-free supernatants were collected and stored at - 20ºC subsequent IAA concentration measurements. For IAA analysis, 100 µl of freshly prepared Salkowski’s reagent (12 mg/ml FeCl_3_ in 7.9 M H_2_SO_4_) (Glickmann and Dessaux, 1995) was added to the same volume of 1/5X diluted samples in flat-bottom 96-wells microplates and incubated for 30 min at room temperature (23-26ºC) in the dark. A_540_ absorbance was read using a microplate reader (iMarkTM, Bio-Rad). IAA was used as standard, although other indole acids including IPyA and indole-lactic acid could technically interfere. Calibration curves were included using 10-100 μM IAA in T-N medium. In these samples, ammonium content was measured using Nessler’s reagents as previously reported (Calatrava et al., 2018).

### Cell quantification by quantitative PCR

The simultaneous quantification of *Chlamydomonas reinhardtii* and *Methylobacterium aquaticum* cell number was inferred by the quantification of the algal- and bacterial-specific single-copy genes centrin and *rpoB* as previously detailed in (Calatrava et al., 2018).

### Statistical analysis

Data represent averages ± Standard Deviation. T-tests were performed using GraphPad Prism 6 with α=0.005, and 3 biological replicates (n=3) unless otherwise indicated in the figure caption.

## Acknowledgements

We thank Isabel M. García Magdaleno from the Department of Mass Spectrometry at the Central Service for Research Support (SCAI) at the University of Cordoba for her technical support in HPLC measurements; Vicente Mariscal and Yongjian Qiu for helpful comments on the manuscript; and María Isabel Macías Gómez for technical support. This study was supported by Ministerio de Ciencias e Innovación (Grant PID2020-118398GB-I00), the European Regional Development Fund program, Junta de Andalucía (BIO-502), the Plan Propio de la Universidad de Córdoba, and the U.E.INTERREG VA POCTEP-055_ALGARED_PLUS5_E. EFYH was supported in part by NSF CAREER grant DEB-1846376.

