## Supplemental Figures and Tables for "Auxin production in the green alga *Chlamydomonas* involves an extracellular L-amino acid oxidase and supports algal-bacterial mutualism with methylobacteria"

**Table S1. Expected α-keto acids produced by LAO1 activity for the 20 proteinogenic amino acids.**

| L-Amino acid | α-Keto acid |
|---|---|
| L-Tryptophan | Indole-3-pyruvic acid* |
| L-Phenylalanine | Phenyl-3-pyruvic acid* |
| L-Methionine | 2-Oxo-4-thiomethylbutanoic acid |
| L-Leucine | α-Ketoisocaproic acid |
| L-Isoleucine | 3-Methyl-2-oxovaleric acid |
| L-Tyrosine | 4-Hydroxyphenylpyruvic acid |
| L-Valine | α-Ketoisovaleric acid |
| L-Glutamine | 2-Keto-glutarate |
| L-Serine | 3-Hydroxypyruvic acid |
| L-Lysine | α-Keto-ε-aminocaproate |
| L-Asparagine | Oxaloacetamide |
| L-Alanine | Pyruvic acid |
| L-Arginine | 2-Oxo-5-guanidinopentanoic acid |
| L-Histidine | Urocanate |
| L-Glutamate | Ketosuccinic acid |
| L-Aspartate | Oxaloacetate |
| L-Threonine | 2-Hydroxy-2-oxobutanoate |
| Glycine | Glyoxylate |
| L-Proline | - |
| L-Cysteine | 3-Mercaptopyruvic acid |

*Auxin precursor

**Table S2. Testing the presence of indole-3-acetic acid (IAA) produced from indole-3-pyruvic acid (IPyA) by incubation with algal cells or hydrogen peroxide.**

| | IAA (mM) |
|---|---|
| IPyA (standard) | N.D. |
| IPyA + *Chlamydomonas* | N.D. |
| IPyA + $H_2O_2$ (without cells) | N.D. |

IPyA (1 mM) was incubated with *Chlamydomonas* wild-type cells ($5x10^6$ cells/ml) or with hydrogen peroxide (1 mM) for 48 h. IAA was analyzed by HPLC in cell-free supernatants. N.D., not detected (i.e., < 0.125 mM)

**Table S3. Ammonium was not detected in the supernatant of *Methylobacterium* spp. incubated with IAA.** Mono-cultures were incubated in N-free media supplemented with 500 µM of IAA for five days. Ammonium was assayed in the cell-free supernatants using Nessler's reagent (see *Methods* section). N.D., not detected.

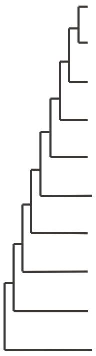

| *Methylobacterium* spp. | Ammonium |
|---|---|
| *Mory* | N.D. |
| *M88A* | N.D. |
| *Morg* | N.D. |
| *M017* | N.D. |
| *Mmar* | N.D. |
| *Mhis* | N.D. |
| *Mnod* | N.D. |
| *Maqu* | N.D. |
| *Maer* | N.D. |
| *Mext* | N.D. |

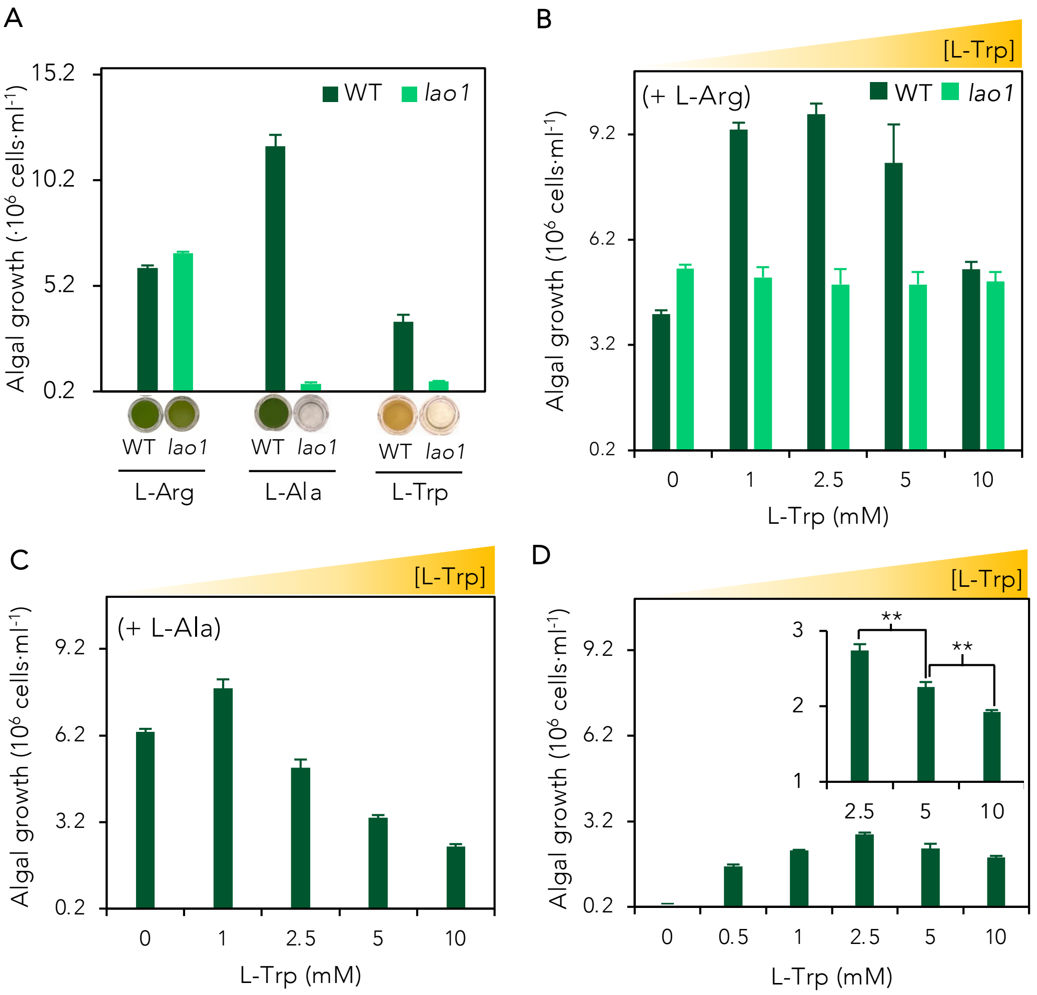


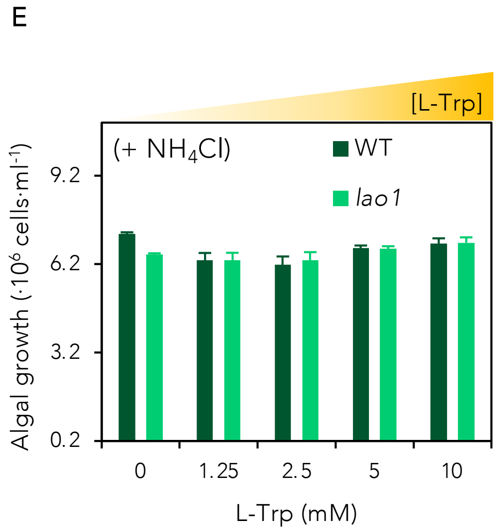


**Figure S1. Impact of L-tryptophan (L-Trp) and LAO1 in *Chlamydomonas* growth on different nitrogen sources**. (*A*) growth on L-arginine (L-Arg), L-alanine (L-Ala) and L-Trp (8 mM) as the sole nitrogen (N) source after 3 (L-Arg and L-Ala) or 7 days (L-Trp). (*B*) wild-type (WT) and *lao1* mutant (*lao1*) cell growth on 4 mM L-Arg supplemented with L-Trp for 3 days. (*C*) WT cell growth on 4 mM L-Ala supplemented with L-Trp for 3 days. (*D*) WT growth on different concentrations of L-Trp as the sole N source for 7 days. Inset shows zoom in. Asterisks indicate statistically significant differences compared to the control (t-test: n=3; α=0.05). (*E*) WT and *lao1* mutant cells grown on ammonium (4 mM) supplemented with L-Trp for 3 days. WT and *lao1* mutant cells were cultured at an initial concentration of $0.2 \times 10^6$ cells/ml. Data shown are of averages of 3 biological replicates and error bars depict standard deviations.

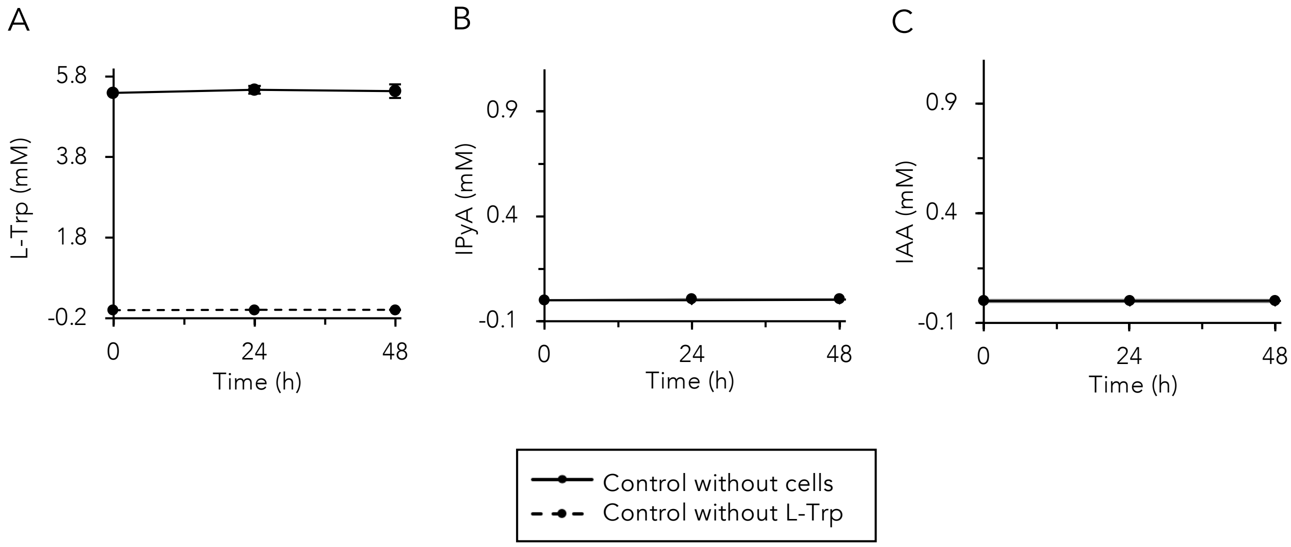


**Figure S2. Controls for *Chlamydomonas* biosynthesis of indole-3-acetic acid (IAA) from L-tryptophan (L-Trp) via indole-3-pyruvic acid (IPyA).** A control without cells (but with L-Trp) and other without L-Trp (but with wild-type cells) were incubated for 48 h in nitrogen-free medium to rule out potential abiotic generation of IPyA or IAA, or biotic but L-Trp-independent production. (*A*) L-Trp, (*B*) IPyA and (*C*) IAA were quantified in the cell-free supernatant/medium using HPLC. Data shown are averages of three biological replicates and error bars depict standard deviations.

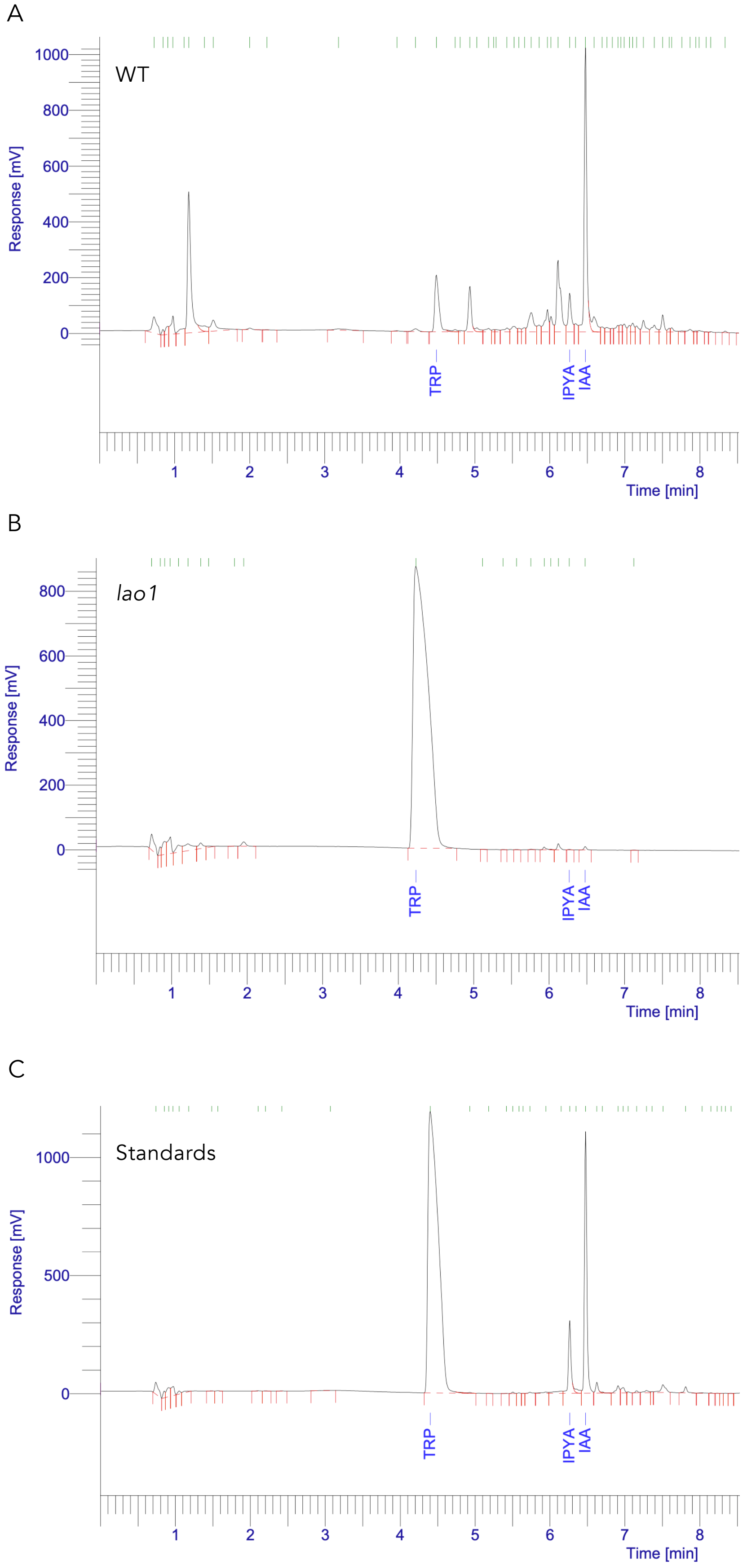


**Figure S3.** HPLC chromatograms for the *Chlamydomonas* WT and *lao1* mutant supernatants corresponding to the results shown in Figure 1. (*A*) Wild-type (WT) and (*B*) *lao1* mutant cells were incubated for 48 h in nitrogen-free medium supplemented with 5 mM L-tryptophan (TRP). Initial cell concentration was $5 \times 10^6$ cells/ml. A representative chromatogram for the three biological replicates is shown. (*C*) Mix of 5 mM TRP, and 1 mM of indole-pyruvic acid (IPYA) and indole-3-acetic acid (IAA) used as a standard.

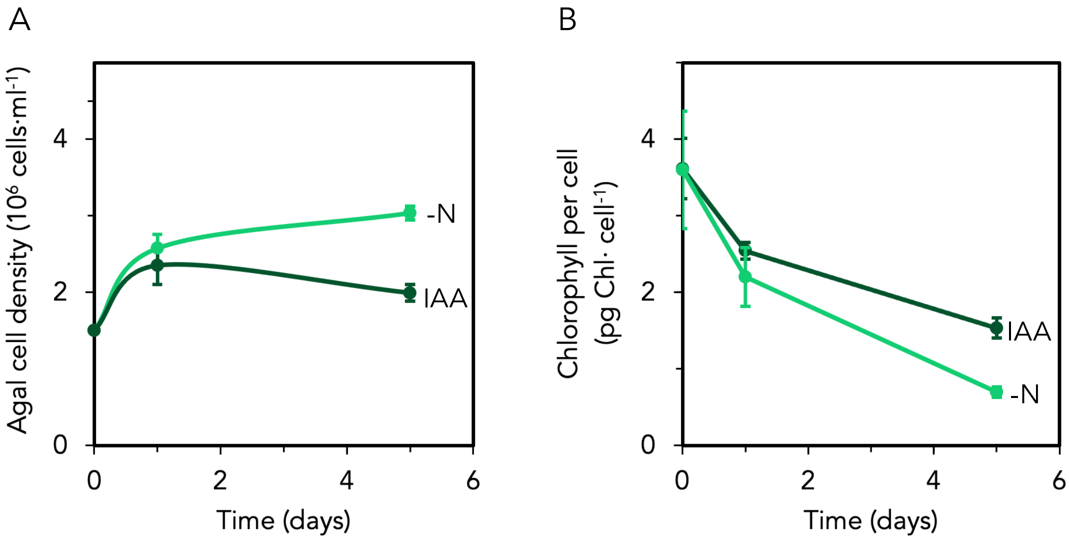


**Fig. S4. Effect of IAA on cell multiplication and chlorophyll content during nitrogen deprivation of *lao1* mutant *Chlamydomonas* cells**. *Lao1* mutant cells were incubated in nitrogen-free media (–N) or supplemented with 500 µM indole-3-acetic acid (IAA). (*A*) The algal cell density and (*B*) chlorophyll content were determined at the at the indicated times. Data show are averages of three biological replicates and error bars depict standard deviations.

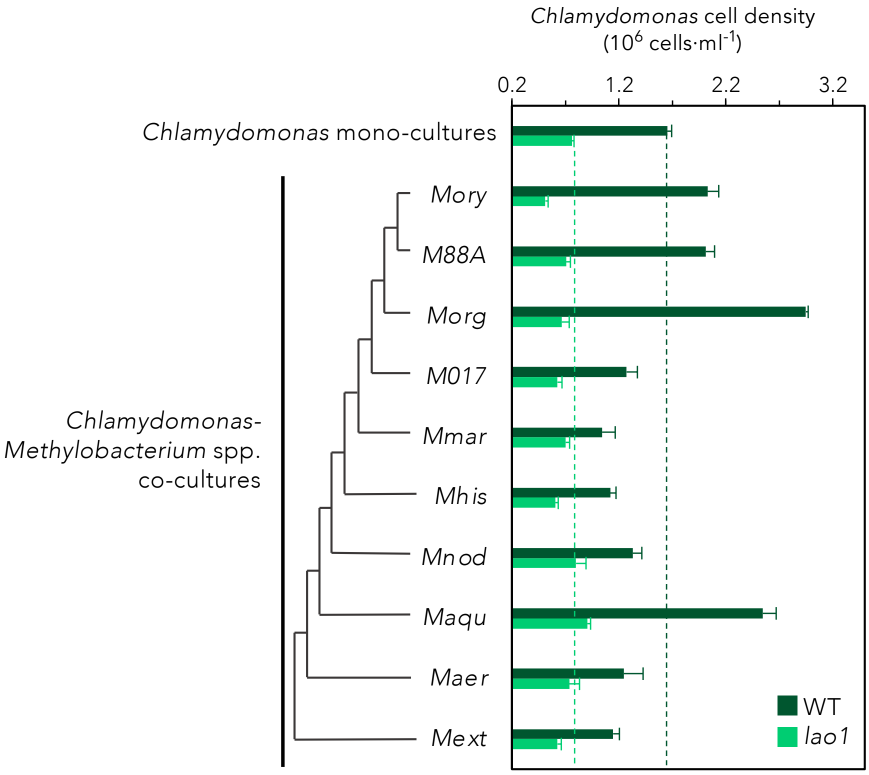


**Figure S5. Algal growth during methylobacterial co-culture on L-tryptophan**. *Chlamydomonas* cell concentration was determined after seven days of growth on 4 mM of L-tryptophan as the sole nitrogen source. Maximum likelihood tree was built using 16S rDNA sequences using MEGAX with default settings. *Mory, Methylobacterium oryzae; M88A, Methylobacterium* sp. 88A*; Morg, M. organophilum; M017, Methylobacterium* sp. M017*; Mmar, M. marchantiae; Mhis, M. hispanicum; Mnod, M. nodulans; Maqu, M. aquaticum; Maer, M. aerolatum; Mext, Methylovorus extorquens (*previously known as *Methylobacterium extorquens)*. WT, wild-type strain; *lao1*, *lao1* mutant strain. Data shown are averages of three biological replicates and error bars depict standard deviations.
